# Inferring learning rules during de novo task learning

**DOI:** 10.1101/2025.09.29.679295

**Authors:** Victor Geadah, Jonathan W. Pillow

**Author notes:** Correspondence: {, }.

## Abstract

Identifying the learning rules that govern behavior is a central problem in neuroscience. While reinforcement learning (RL) offers a unifying theoretical framework, most empirical studies of animal learning behavior have focused on non-stationary environments (e.g. changing reward probabilities in a known task), as opposed to acquiring an entirely new task from scratch. Here we introduce a statistical framework to infer reinforcement learning rules directly from single-animal behavior. Applied to mice learning a perceptual decision-making task, our approach reveals that policy-gradient–like rules capture de novo task learning better than classical temporal-difference algorithms. By fitting flexible parametric learning rules, we uncover systematic deviations from standard RL models, including side-specific learning rates and negative reward baselines. Together, these parameters account for side-biased learning, as well as forgetting and consecutive errors due to aversive responses to incorrect trials. Extending the framework with latent, dynamic learning rates further reveals that animals adapt their learning rates over training and across curricula. These results provide a statistical account of how animals learn from scratch and highlight key departures from classical reinforcement learning algorithms.

## 1 Introduction

Training animals to perform novel tasks is a cornerstone of neuroscience experiments, yet the learning process itself remains undercharacterized. The training process is laborious, taking likely weeks of shaping and feedback before reaching satisfactory performance and requiring significant time and lab resources [1, 2]. Despite this centrality, the dominant focus is on the resulting trained behavior, and we still have a limited understanding of how animals learn to perform a task *de novo*.

Reinforcement Learning (RL; [3, 4]) provides a normative framework to understand how tasks can be solved and acquired from reward feedback. From early models of classical conditioning [5, 6] to seminal neuroscientific accounts of error-driven learning [7], RL-based formalisms have provided unprecedented insights into theories of optimal behavior—how an agent *should* act. Yet much of the animal RL literature studies behavioral *adaptation in trained animals*, using using bandit-style tasks with nonstationary or hidden reward contingencies [8–12]. Indeed, paradigms such as probabilistic reversal learning [10, 11] and drifting multi-armed bandits [12] now dominate, with behavior commonly modeled by trial-to-trial RL or probabilistic inference frameworks. This literature has mapped robust neural correlates of value updating and exploration–exploitation in already-trained animals. By design, these settings offer a more controlled way to vary only specific modalities of the environment (predominantly reward contingencies), isolating variables and narrowing the search of algorithms and models [13], but they do not capture the early-training dynamics of de novo learning where assumptions of stationarity and near-optimality fail. This early learning, “training data”, is in contrast often discarded and largely ignored.

In practice, animals often deviate from experimenter-based notions of optimality: they have biases, lapses, and other idiosyncratic tendencies. This challenge is particularly stark during de novo learning. Even in simplified two-alternate force choice task with fixed reward contingencies, mice show significant inter-animal heterogeneity in learning dynamics [14, 15], even despite highly standardized training protocols [1]. Learning is further governed by internal states, such as arousal, motivation, and engagement, which introduce structured, nonstationary fluctuations in behavior [16–18]. Any account of de novo learning must therefore allow for structured idiosyncrasies, context dependence, and latent nonstationarity.

Probabilistic and statistical methods let us approach this variability by drawing a more data-driven account of the learning process. They excel at giving us insights into behavior, but typically do so with descriptive methods [16–18] that lack a mechanistic model for the internal states—a model of *how*, less so *why*, they evolve. Marrying probabilistic techniques within normative models is a promising direction to tackle empirical variability itself, for instance by introducing stochasticity itself in the value [19] and perceived reward [20] learning dynamics. While some previous work has laid the modeling and theoretical grounds for inferring learning rules directly from behavioral data [21–23], it hasn’t been brought to scale and to single-animal inference to aid our understanding of de novo learning.

To overcome these limitations, we derived a framework to infer, at the single-animal level, empirical reinforcement learning rules driving learning dynamics during de novo task learning (§2.1). We used this approach to analyze mice learning to perform, from scratch, the International Brain Laboratory (IBL) binary perceptual decision-making task. We inferred, per animal, precise forms of reinforcement learning rules employed (§2.2) and provided decompositions for quantification and predictions of learning dynamics (§2.3). Finally, we identified dynamic modulation of learning rules by latent dynamic learning rates (§2.4) in two IBL training curricula variants.

## 2 Results

In this work, we investigate perceptual decision-making data of mice learning to perform a two-alternative force choice task, collected as part of the International Brain Laboratory (IBL). The task consists of moving a wheel to align a visual Gabor grating (Fig. 1**a**) in the center, after which the water-deprived animals receive a sugar-water reward on correct trials. After a habituation period, mice are trained to perform the task at varying contrast levels but with equal probability of the stimuli appearing on either side, before moving to a block-trial structure with alternating 80/20% probabilities (Fig. 1**b**). We selected for mice that attained the performance level required to move to the block-switching component of the task, but only considered the training days portion of the dataset. In all, our primary dataset consists of choice data of *N* = 31 mice, trained for a median of 22 days totaling 12708 trials, each (Fig. 1**c**).

**Figure 1:**
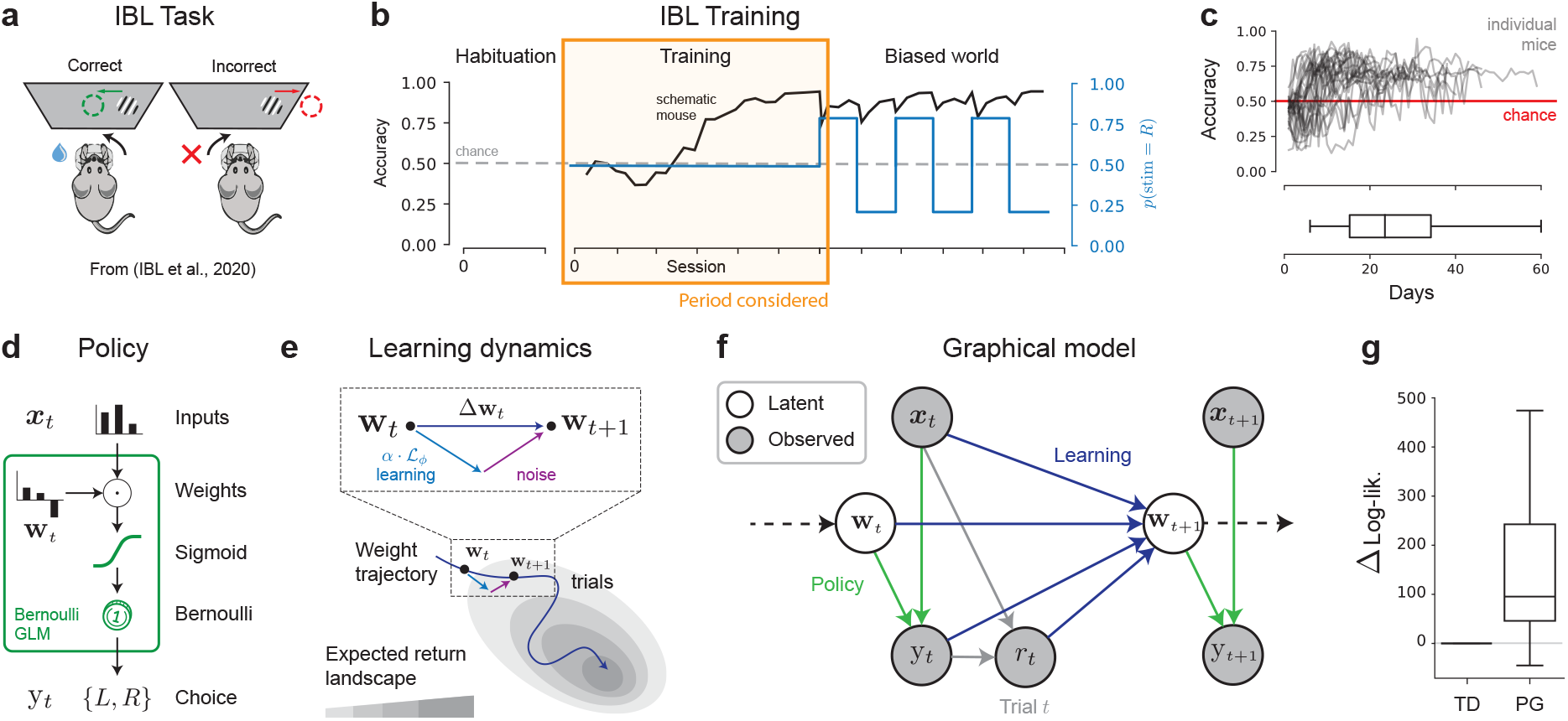
Learning data and modeling. (**a**) Experimental perceptual decision-making set-up from the IBL. (**b**) The IBL mouse training paradigm consists of three consecutive protocols: (1) habituation to the rig and stimulus-action association, (2) “training” protocol, where mice go from chance-level to satisfactory performance, and (3) side-biased reward probabilities in alternating block structure. In this work, we focus on the second protocol. (**c**) Accuracy (fraction of trials rewarded) per session of training for the *N* = 31 mice considered (top), along with distribution of total days of training (bottom). (**d**) We model action selection (policy) with a Bernoulli Generalized Linear Model. (**e**) Learning is modeled as the evolution of the weights **w**_*t*_ to maximize the expected return according to a learning rule ℒ, along with additive noise on each update. (**f**) Combining the policy and learning dynamics models yields the full probabilistic graphical model for the data. From the perspective of the experimenter, the inputs, choices and rewards are observed, and the weights **w**_*t*_ driving the decisions are latent. (**g**) Difference in log-likelihood between the GLM with policy gradient (PG) learning dynamics and an action-value temporal-difference (TD) learning model. Box plot indicates distribution over animals. Higher is better.

### 2.1 Probabilistic and reinforcement learning model of animal learning

Following previous work, we model the animal’s decisions with a Bernoulli Generalized Linear Model (GLM, see Fig. 1**d**), where the probability that an animal makes decision y_*t*_ ∈ {*L, R*} at trial *t* ∈ {1, …, *T* } given a vector of task covariates ***x***_*t*_ ∈ ℝ^*M*^is expressed with the logistic link function

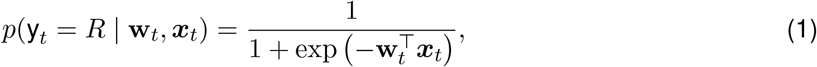

and *p*(y_*t*_ = *L* | **w**_*t*_, ***x***_*t*_) = 1 − *p*(y_*t*_ = ∈ *R* | **w**_*t*_, ***x***_*t*_), with weights **w**_*t*_ ∈ ℝ^*M*^describing the influence of each task covariate. These covariates ***x***_*t*_ ∈ ℝ^*M*^ include task-relevant features, such as the stimulus intensity, and other covariates that might influence an animal’s choice, such as previous rewarded side or previous choice. This decision-making model *p*(y_*t*_ | **w**_*t*_, ***x***_*t*_) acts as our *policy*, mapping stimuli (states) to decisions (actions), with weights **w**_*t*_ acting as policy parameters.

Mice learn from decisions and rewards. We cast learning as updating the policy weights **w**_*t*_ over trials, evolving according to a learning rule ℒ_*ϕ*_

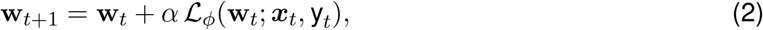

with learning rate *α >* 0. The learning rule ℒ_*ϕ*_ is a function of the previous weights, the previous decision, and the previous stimuli, with parameters *ϕ*. We will use reinforcement learning (RL) to model ℒ_*ϕ*_, and devising or inferring its precise form will be the topic of the sections to follow.

#### 2.1.1 Model fitting and inference

Our probabilistic model (Fig. 1**f**) of decision-making learning dynamics can be described by the policy in eq. (1) and the weights evolving with mean from eq. 2 and additive Gaussian noise

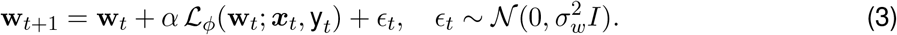

See Fig. 1**f** for a schematic of the resulting learning model, with parameters *θ* = {*α, σ*_w_, *ϕ*}. This adds noise in the dynamics of the weights, which is in contrast to the more classically considered deterministic models of learning. The resulting model is fully probabilistic, allowing for flexible inference and to capture individual variability, in line with recent stochastic models of learning dynamics [19].

We evaluate model performance with the marginal log-likelihood of the data

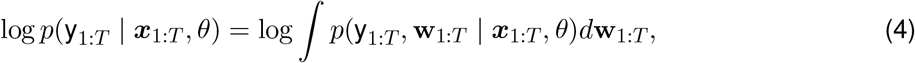

which is the probability, as a function of *θ*, of an observed behavioral sequence y_1:*T*_ given a sequence of regressors ***x***_1:*T*_ and marginalizing over possible weight trajectories **w**_1:*T*_ under our learning rule prior in eq. (3) of parameters *θ*. This marginalization is usually intractable, and here we resort to particle filtering algorithms [24, 25] that estimate this integral from samples sequentially updated from the learning dynamics model. We perform maximum likelihood estimation (MLE) for the parameters *θ* by maximizing the log-lik. with gradient ascent. As a form of cross-validation for fitting to single-trajectories, 10% of the trials were considered as un-observed in the training set—every metric of log-lik. reported in the text refers to the *full* trajectory log-lik., including these held-out trials. The filtering log-likelihoods on the held-out trials are provided in Appendix Fig. 9. Finally, we report posterior error bars and uncertainty for the parameters, obtained with Metropolis Hastings. For more details on our implementation, training algorithm and posterior sampling, see Appendix §C.1.

#### 2.1.2 Focus on policy gradient over temporal-difference learning

Accounts of *learning* with reinforcement learning focus on studying how agents update their understanding of their environment, typically with the goal to maximize the *expected-return*

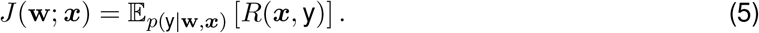

under the policy (Fig. 1**e**). Here, *R*(*s, y*) is the scalar reward obtained from making a decision *y* when presented the stimulus intensity *s* (or equivalently ***x***), taking values *r*_*t*_ on trial *t* of *r*^∗^ *>* 0 if correct and 0 otherwise. There is no discount factor due to our simple one-time-step decision process, making the above equivalent to the *state-value function* [3, eq. (3.12)].

Our decision-making model is a parametrized policy with parameters **w**, making **policy-gradient** (PG) [3, 26, Chapter 13] a natural choice for modeling ℒ. Previously used in modeling learning dynamics [21, 22], PG methods cast learning as performing gradient ascent of the expected-return objective, with 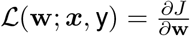. The gradient of this expectation is typically not available in closed-form and a popular approximation is given by the REINFORCE [27] algorithm, which leverages a proportional quantity to the true gradient and estimates it with Monte Carlo samples to arrive at

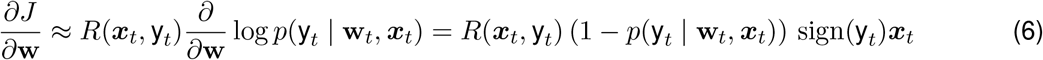

with the second inequality being the resulting expression in our setting [22].

Action-value methods are a contrasting approach to maximizing the expected return in a model-free way [3, 9]. They posit that the agent learns to construct values for the different states encountered in its environment, and uses said values to make decisions. They are central to work focused on hidden or varying reward contingencies, and less but nonetheless still considered within perceptual decision-making tasks [28]. We considered a particular standard model, the temporal difference RL (TDRL) model from Ref. [28], in which the agent (1) uses Bayesian inference to estimate the true stimulus *s*_*t*_ from partial observations, the percept 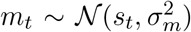, (2) uses this belief about the stimuli to guide the decision-making based on choice values ***V***_*t*_ = [*V*_*L,t*_, *V*_*R,t*_] at trial *t*, and (3) updates these values by temporal difference (TD) learning. This model thus covers both partial and fully observed MDP treatments of behavior, as well as stochasticity in decision-making through a temperature parameter (see Appendix B.1 for a full description). We found that PG, with matched number of weights, attained significantly higher marginal log-likelihood that TDRL on the data (Fig. 1**g**). Thus, we elected to consider policy-gradient learning methods with Bernoulli GLM policies as our focus of study for the reinforcement learning dynamics.

As support for this direction, we investigated key commonalities and differences between the TDRL and PG formulations (App. B.2). Focusing first on the policy, marginalizing over the noisy percept shows that the TDRL choice rule can be written as

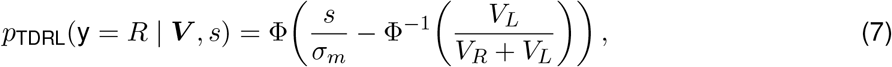

where Φ is the standard normal CDF—a standard signal detection theory argument [29, 30] (derivations in App. B.2.1). Thus, the TDRL policy can be cast as a Bernoulli GLM with probit (or logit) link, making the TDRL and PG models policy-equivalent. As for the learning dynamics, the two approaches share confidence-guided [28] weight updates (App. B.2.2) but differ in how noise enters the update. Averaging over percepts, the TDRL value update is approximately

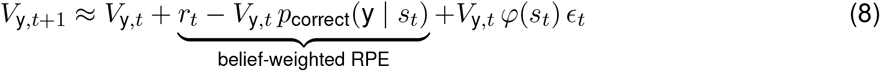

with *ϵ*_*t*_ ∼ 𝒩 (0, 1), belief term 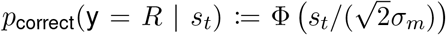, the two-channel success probability [29], and *φ* the PDF of 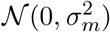 (App. B.2.3). Thus perceptual uncertainty induces an additive fluctuation whose magnitude grows with larger current values and lower-contrast stimuli. By contrast, in our PG model (eq. 3) we assume a more flexible, trial-independent additive noise process [16, 19, 22].

### 2.2 Learning rule inference: mice forget and learn sides at different rates

We sought to infer a flexible form of policy-gradient dynamics for each animal. First, we expanded on our set of regressors and consider ***x***_*t*_ ∈ R^5^ to consist of *M* = 5 elements: (1) a bias term (always equal to 1), (2) the stimulus contrast level on the left side (0 if absent), (3) the stimulus contrast on the right side, (4) the previous choice y_*t*−1_, and (5) the previous correct side *s*_*t*−1_ ∈ {*L, R*}. Then, we considered a flexible parametric expression for the policy gradient update where every parameter, such as the learning rate or additive noise, is learned to be specific to each regressor.

Concretely, we considered the parametric policy gradient update

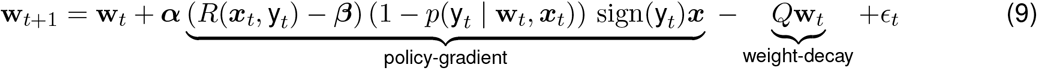

with additive noise 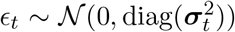. We introduced a vector of learning rates ***α*** ∈ ℝ^*M*^ along with per-regressor noise scales ***σ***_*t*_ ∈ ℝ^*M*^, taking values in {***σ, σ***_*day*_}. We also included a weight decay of the form −*Q***w** with the diagonal matrix *Q* ∈ ℝ^*M×M*^, which could account for forgetting. Finally, we used the REINFORCE form to the policy gradient update, which accommodates a baseline parameter ***β*** ∈ ℝ^*M*^ to the reward. Every vector multiplication in eq. (9) is taken to be point-wise. We refer to this vectorized model as **vector PG**, with parameter set *θ* ={ ***α, σ, σ***_*day*_, ***β***, *Q*}. As baselines, we used a **scalar PG** parametrization, with the same but scalar parameters, and the PsyTrack model from [16] which does not account for any learning model (i.e. setting *α* = 0, see Appendix §B.1).

We found the inferred learning rule to differ from more widely used variants. A standard usage of REINFORCE for policy-gradient learning does not account for a baseline (***β*** = 0), in which case learning occurs only on correct trials (Fig. 2**a**-(left)). A more broadly used variant and natural choice is a positive baseline corresponding to the expected return for each regressor (Fig. 2**a**-(right), also akin to Actor-Critic [3] methods), which leads to learning governed by reward prediction errors [5], learning more on incorrect than correct trials. When fitting our vectorized parametric form to the data, we found significant diversity across animals, and namely two primary distinctions from classical rules: (1) different learning rates per side, and (2) a *negative* baseline for reward (Fig. 2**b**-**c**). This vectorized learning rule model achieved higher marginal log-likelihood (Fig. 2**g**) on whole trajectories, which we recall includes test trials excluded from training.

**Figure 2:**
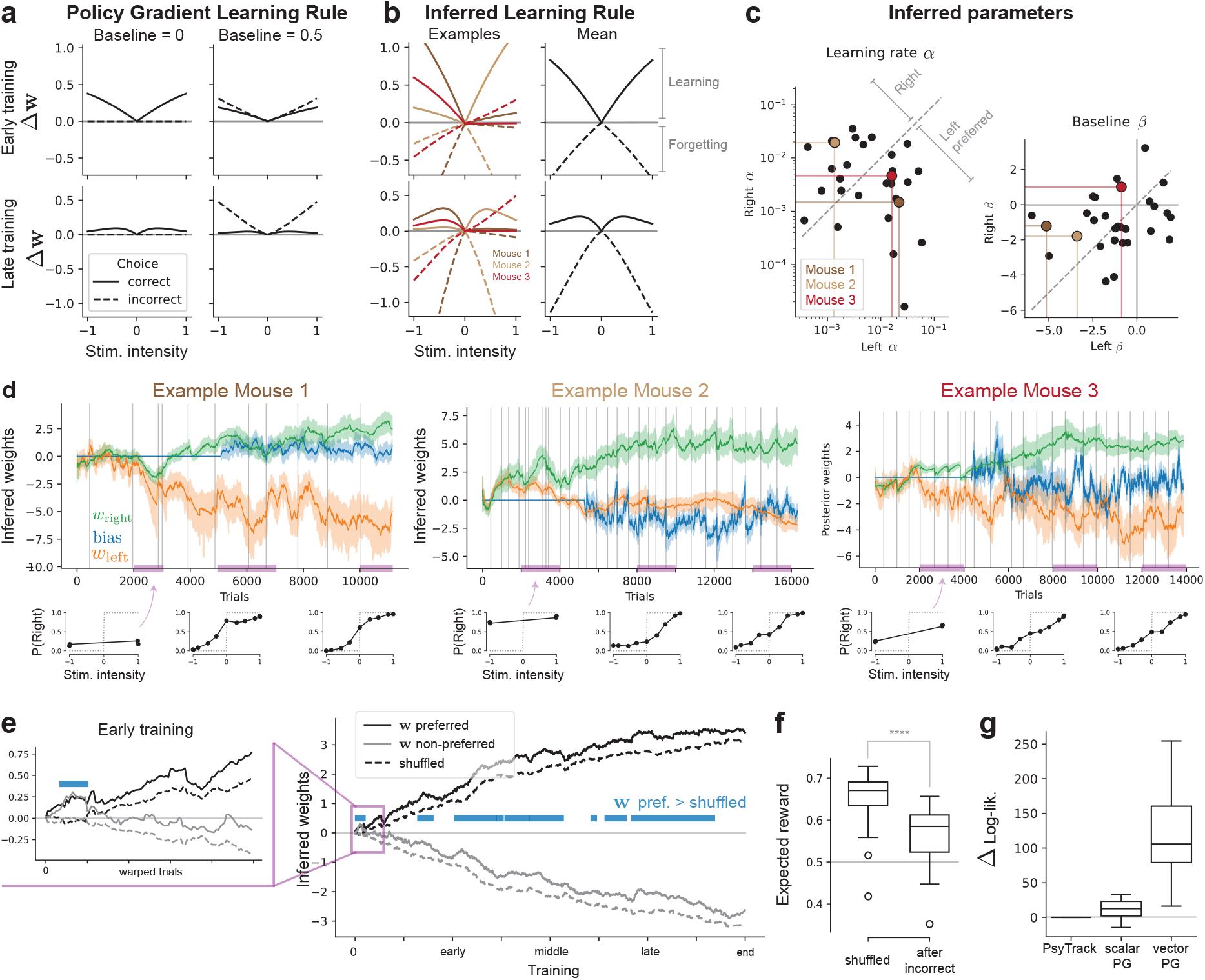
Parametric learning rule inference. (**a**) Classical policy gradient learning rule updates Δ**w**, for early (top) and late training (bottom), with *β* = 0 (left) and *β* = 0.5 (right). (**b**) Inferred learning rule displays asymmetric learning and a negative baseline for reward, with negative updates on incorrect trials. Learning rules of example mice (left) and mean over mice (right) are shown. (**c**) Inferred parameters per mice in parameter space. Same example mice as in (**b**) are highlighted. (**d**) Weight posterior ŵ_1:*T*_ trajectories for the three example mice of (**b**-**c**) (top) along with corresponding data psychometric curves (bottom) highlight. First two highlight side-preferences in learning, last example shows symmetric learning. Error bars indicate posterior standard deviation (±1 SD) about the mean, faint vertical gray lines indicate training days. (**e**) Average of posterior mean over animals, sorted by preferred and nonpreferred weights depending on the learning rates (see text). Insert on the left shows early learning. Blue bars indicate when the preferred weight is higher than the shuffled control (*p <* 0.05, one-sided one-sample *t*-test). (**f**) The expected reward after an incorrect trial (𝔼 [*r*_*t*+1_ | *r*_*t*_ = 0]) is lower than the shuffled control (𝔼 [*r*_*t*_]). Box plot indicates distribution over animals. (**g**) Difference in log-lik. with respect to PsyTrack as baseline (higher is better), plotting the box plot of the distribution over animals.

First, multiple studies in IBL mice have identified side-preferential learning [14–16]. We first examined this behavior directly through the inferred learning rate parameters *α*_left_ and *α*_right_ across animals (Fig. 2**c**). We compared the absolute difference between the two side-specific learning rates from zero using a one-sided one-sample *t*-test, which revealed a highly significant effect (*p* ≪ 0.001), indicating that mice do indeed learn one side before the other (see first and second example trajectories in Fig. 2**d**). A paired *t*-test of left versus right learning rates showed no significant difference (*p >* 0.05), suggesting that the side learned first was distributed symmetrically between left and right across animals. Second, equipped with these single-animal estimates of the learning rates, we can examine the inferred learning dynamics. We re-labeled left and right weights {**w**_*t*,left_, **w**_*t*,right_} to {**w**_*t*,preferred_, **w**_*t*,non−preferred_}, where the preferred side is defined by the higher learning rate *α* between left and right. We plot in Fig. 2**e** the resulting posterior trajectory for these two preference-based weights, averaged over animals and compared against a control where the parameters and corresponding animals are shuffled. We found that the side-preferred weight is often higher than the shuffled control, attesting to side-preferential overall learning in these mice.

Second, mice displayed a negative baseline for reward (*p <* 0.001, one-sided *t*-test on ***β*** over animals). In terms of learning updates, this negative baseline causes a *negative* weight update on incorrect trials (Fig. 2**b**). This predicts that mice would get worse after incorrect trials, and furthermore in early training this can cause a stimulus weight to increase in the wrong direction. Looking at the preferred-sorted weight early in training (Fig. 2**e**, left inset), we found biased learning with both weights deviating together in the preferred direction. This further compounds the biased learning mentioned above. We further validated the ramifications of this negative baseline by analyzing the data directly (Fig. 2**f**)—we found that the expected reward following an incorrect trial was considerably lower than the shuffled expected reward (*p* ≪ 0.001). We found that removing the weight-decay term *Q* did not significantly impact marginal log-likelihood performance (*p >* 0.05, one-sample *t*-test, over subjects), hence the forgetting phenomena was primarily accounted for by the negative baseline in our model formulation.

### 2.3 Predicting learning dynamics; learning and noise decompositions

The inferred weight trajectories capture simultaneously the learning of the animal and its stochasticity. Our learning dynamics models (Fig. 1**e**) posit that learning update can be decomposed into a learning term, *α* ℒ_*ϕ*_, and a noise term *ϵ*_*t*_. We sought to leverage this modeling to establish a decomposition of entire posterior trajectories ŵ_1:*T*_ into a *learning component* and a *noise component* (Fig. 3**a**).

**Figure 3:**
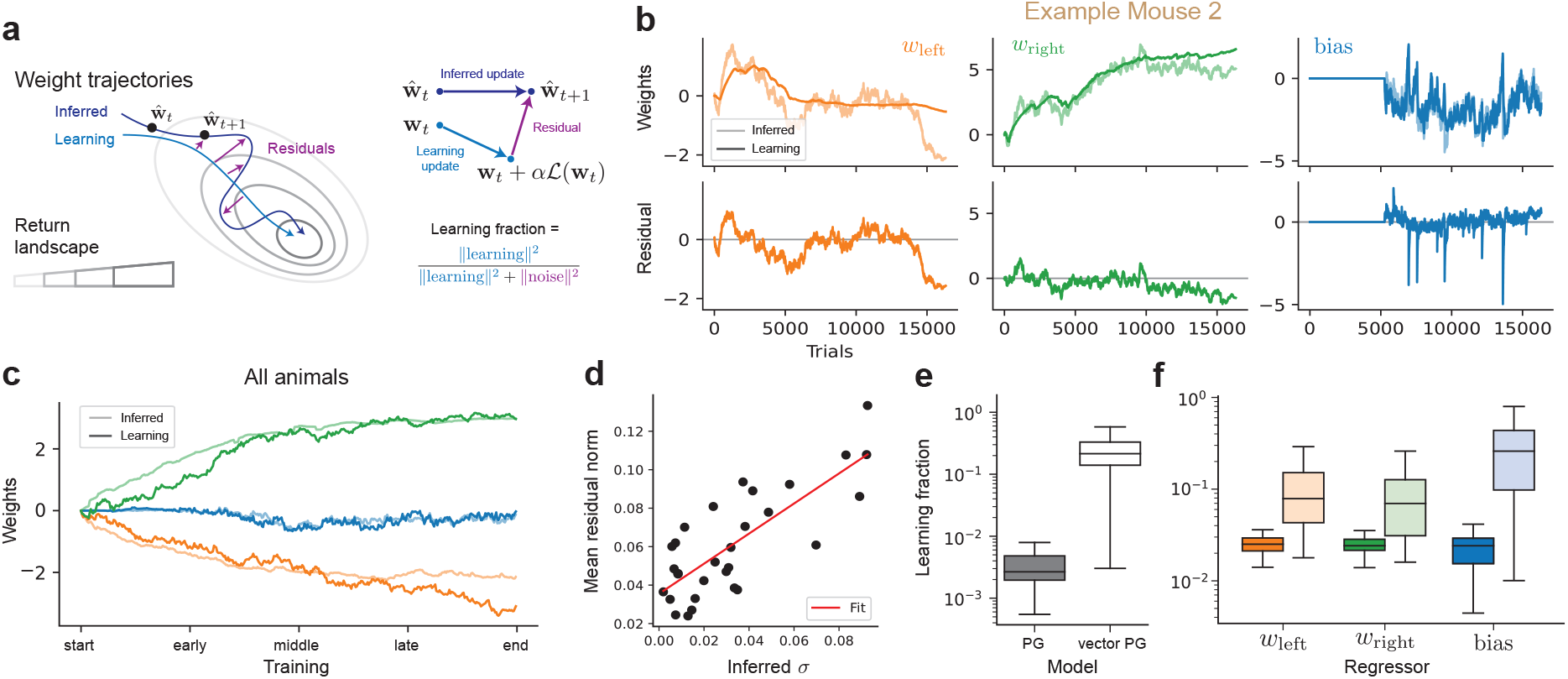
Decomposing dynamics into learning and noise. (**a**) (right) The *learning* component from our model is defined as the deterministic mean of the predictive, generative process. (left) The residuals between the learning trajectory and the posterior mean (“inferred”) define the noise component, in turn used to define the “learning fraction”. (**b**) (top) Overlay of the inferred weights and the learning trajectory for the vector PG model for the example mouse #2 from Fig. 2, per weight, along with (bottom) residual, obtained as the difference between the inferred and learning trajectories. (**c**) Average over all mice (*N* = 31) of the inferred and learning trajectories largely overlap. (**d**) Average over trials of the residual norm and inferred noise scale *σ* = ∥***σ***∥ per animal, showing a high correlation (*R*^2^ = 0.62, *ρ* = 0.79). (**e**) Learning fractions per model, with higher values (up to 1) indicating better alignment between the posterior and the learning model. (**f**) Learning fractions per regressor for the scalar (dark) and vector (light) PG models.

Our approach consisted of defining the learning component of any individual decision-making trajectories as the predictive mean under our model (Fig. 3**a**-left). This is obtained by updating the weights **w**_*t*_ using the learning rule in eq. 2 with the animal’s decisions y_*t*_, without any noise. The *noise* component of the dynamics is then taken to be the residual between the posterior mean (the “inferred” trajectory) and learning component (Fig. 3**a**). We plot in Fig. 3**b** the resulting learning and inferred trajectories overlaid for an example mouse, reflecting a high-degree of alignment and capturing some of the idiosyncrasies of the learning dynamics of this particular mouse. The learning decompositions for all three example mice can be found in Appendix C.2, and we plot in Fig. 3**c** the learning and posterior trajectories, averaging over the *N* = 31 animals considered. Finally, we found (Fig. 3**d**) this definition of the noise component to match the inferred parameter values of noise scale *σ* per animal with high correlation (*ρ* = 0.79), providing support for this learning decomposition as consistent with our modeling.

We found the learning component to recapitulate the key learning weight dynamics of the inferred posterior mean, such as the primary side dominance and even the trial-to-trial “noisy” fluctuation in weights and bias. To quantify how much the learning component accounts for the inferred weights trajectory on every trial we considered the *learning fraction* (Fig. 3**a**-right), defined as the ratio between the norm of the learning update *α* · ℒ_*ϕ*_(**w**_*t*_; y_*t*_, ***x***_*t*_) and the norm of the residual noise with the inferred weight update ŵ _*t*+1_ − ŵ _*t*_ (Fig. 3**e**-left), averaged over all trials. For reference, a noise-only model like PsyTrack [16] has a learning fraction of strictly 0, and a deterministic learning-only model 1. We include in

Appendix C.2 alternate metrics, such as projection fraction, cosine similarity and the one considered in previous work [22]—we found no qualitative difference. We found the vector PG accounted for significantly more of the learning component (Fig.3**e**) than the scalar version, as a distribution over individual animals. This learning fraction was consistently higher across regressors considered, with the largest difference being for the bias (Fig. 3**f**). This supports our inferred model as an appropriate mechanistic model, able to generate and recapitulate the learning dynamics observed empirically.

### 2.4 Dynamic learning rate

So far, the learning rate *α* governing the learning dynamics has been static, optimized as a hyper-parameter with MLE. This assumes the learning rule is engaged with equally over learning; a restrictive assumption richly studied [31], with further recent work [17, 18] pointing to the contrary through nonstationarity in internal states over learning. To provide a dynamic generalization of our learning model, we placed a random-walk prior on the learning rate as

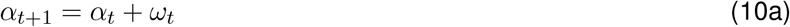

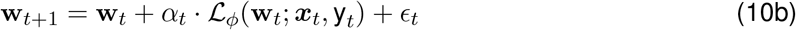

with 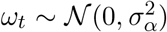 and initial condition 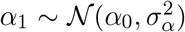, with **w**_1_ and 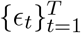 sampled as in the static case. The learning rule ℒ_*ϕ*_ is taken to be exactly the vector PG update in (9), including a global learning rate parameter ***α*** dynamically scaled by the latent variable *α*_*t*_—this **dynamic PG** model of eqs. 10 thus encompasses strictly the vector PG model, with additional {*α*_0_, *σ*_*α*_} scalar parameters. The learning rate is now a dynamic latent variable that is fully integrated out (alongside the weights) in computing the marginal log-likelihood, and for which we can compute posterior trajectories over trials *t*. This learning rate acts as a latent gating on the learning signal, dictating how much an animal engages with a certain learning rule on trial *t*.

We found mice to show a dynamic learning rate, which in turn helped predicting the learning component. First, we plot in Fig. 4**a** the difference log 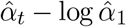 over learning for the per-animal posterior learning rate 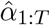 ; since the dynamical learning rate 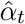 scales a static parameter ***α***, we use the difference in log values which is invariant to such scaling. We found highly dynamic learning rate trajectories in the inferred posterior mean for 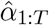, differing across individuals (Fig. 4**a**). Now, we can use this inferred learning rates to refine our prediction the learning component (§2.3) of the trajectory, using the inferred 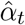 at every time step *t* of the predictive trajectory. We found this to help the learning component more closely match the inferred trajectory (Fig. 4**b**), and generally lead to an increase in the learning fraction (Fig. 4**c**) over animals. The dynamic PG model did provide an increase in marginal log-likelihood over animals (Fig. 4**d**), which includes the held-out test trials attesting to an increase in quality of fit.

**Figure 4:**
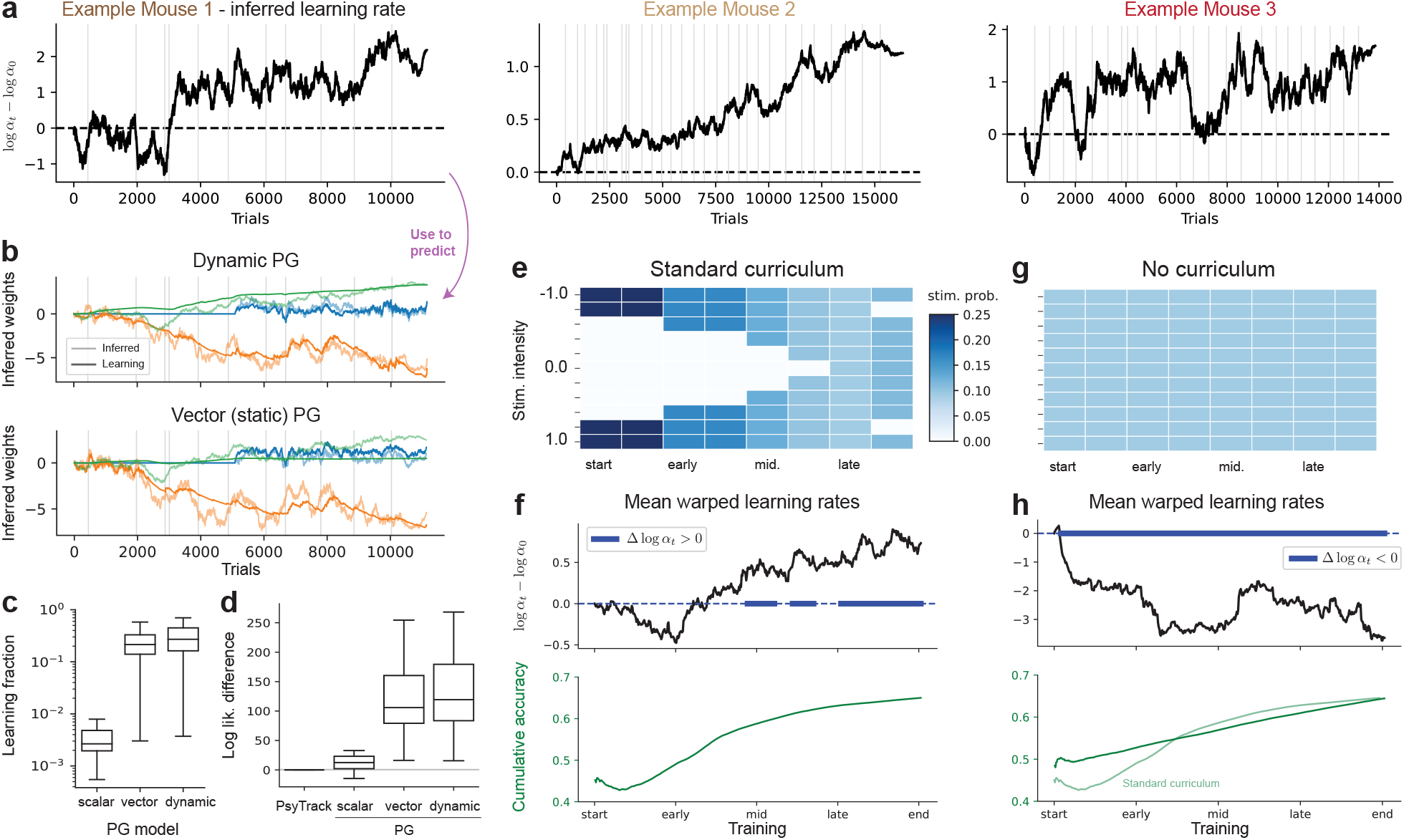
Dynamic learning rate reinforcement learning. The dynamic PG model posits that learning is governed with a dynamic learning rate. (**a**) Inferred (posterior mean) dynamic learning rates for the three example animals considered in Fig. 2-3. Plotted as the deviation from the initial learning rate log *α*_0_ in log-scale. (**b**) We use the inferred learning rate trajectory to predict the learning component, overlaid with the posterior mean for the weights (top), for example mouse #1. Learning component for the vector PG model (bottom) in contrast did not fully capture both stimulus weight dynamics. (**c**) Fraction of learning for the vector and dynamic PG models, with boxplots indicating distributions over animals. (**d**) Difference in log-lik. with PsyTrack per model (higher is better). (**e**) Schematic of the stimulus probability sequence for the standard curriculum used in IBL training. (**f**) Mean over animals of the learning rates difference over training (top), as well as the correlated regressor of cumulative accuracy (green, *ρ* = 0.92), all trial-warped and averaged over animals. Shaded blue bars around Δ log *α*_*t*_ = 0 indicate where the warped learning rates are different (*p* ≤ 0.05, two-sided one-sample *t*-test, across subjects) from 0. (**g**) We investigated a second dataset [15] without any training curriculum, in which all stimuli are presented with equal probability over training. (**h**) Learning rates and cumulative (c.f. (**f**), overlaid in light colored) for the no-curriculum training variant.

Second, we investigated global tendencies in learning rate dynamics. For the standard IBL training curriculum considered so far, which introduces lower contrast stimuli gradually (depicted in Fig. 4**e**), we saw a lot of fluctuations in the average over animals before eventually increasing over the training days (Fig. 4**f**, one-sided, one-sample *t*-test over subjects for log *α*_*t*_ *>* log *α*_0_ at every time-step *t*). We found this average learning rate trajectory to be highly correlated (coefficient of *ρ* = 0.92) with the average cumulative accuracy (Fig. 4**f**-bottom). To elucidate and verify this correlation, we investigated an alternative training protocol with *no* curriculum, in which all the stimuli are presented with equal probability (Fig. 4**g**) from Ref. [15]. In this protocol, we found the learning rate to sharply *decrease* over training (Fig. 4**h**-bottom, *p <* 0.05, log *α*_*t*_ *<* log *α*_0_), with a negative and much weaker correlation to the accuracy (*ρ* = −0.52). Broadly, these results attest to a dynamic learning rate with dynamics dependent on task curricula.

Finally, an alternative would be to consider a dynamic *baseline* for reward. The resulting model could capture learning dynamics of Actor-Critic models, in which case the baseline would evolve from 0 to the expected return per regressor. We found that placing a similar random-walk prior on a time-varying vector of baselines ***β***_*t*_ ∈ ℝ^*M*^ did not significantly outperform the static model nor the dynamic learning rate model (Appendix Fig. 10).

## 3 Discussion

Our results introduce a single-animal framework for uncovering learning rules directly from behavior during de novo task acquisition, revealing rich heterogeneity across mice and marked departures from canonical RL models. By fitting policy-gradient (PG) learning rules to single-animal trajectories, we recover interpretable components (weights, baselines, and time-varying learning rates) that explain learning dynamics without presupposing a fixed, shared strategy. This approach complements recent standardization efforts that enable rigorous cross-lab behavioral comparisons in mice, while focusing on the individualized (and variable) paths by which animals acquire task structure.

We start by expanding on our focus on policy gradient learning rules over temporal-difference (TD) alternatives. We showed (App. B.2.1) that the TDRL policy adopted for the IBL task is reparameterizable as a Bernoulli GLM, so policies alone cannot distinguish the TD from PG formulations considered. Instead, the models differ primarily in their latent update rules. Empirically, we found the simple variants of PG consistently fit better than similarly simplified TD on the same animals (Fig. 1). This accords with emerging evidence that gradient-based credit assignment captures individual learning trajectories in rodents [14], all the while remaining consistent (as we show in Appendix B.2.2)) with behavioral work showing confidence-weighted updating in perceptual decisions [28]. A further diagnostic is the inferred baseline: we obtain a robustly negative baseline (Fig. 2), whereas a positive baseline would better match a classic reward-prediction-error (RPE) signature. In PG, the baseline is a variance-reduction [3, 27] device whose magnitude (and sign) do not alter the expected gradient, making PG less constrained—and, here, more consistent—with the data.

Across animals, we observed learning-rate dynamics (Fig. 4) that are far from constant: learning rates rise and fall over training, even despite a stationary task generative process (no formal curriculum), suggesting that animals *perceive* changes in uncertainty/volatility or adjust step sizes with engagement, arousal, or surprise. These single-animal dynamics echo normative predictions [31] that volatility (such as the one encountered in the standard IBL curriculum) should increase learning rates, while pure stochasticity (e.g., closer to the “no curriculum” variant) should decrease them, and dovetail with observations that neuromodulatory systems adapt the effective rate of learning [6, 31–33]. Together, these findings motivate explicit modeling of latent learning rates as a function of belief dynamics and internal state, not just task statistics.

Our work relates to various notions of cost function inference. A relevant and related class of approaches is *inverse RL* (resp. inverse Optimal Control) [20, 34, 35], which seeks to identify the reward (resp. cost) function used by an agent directly from recordings of its behavior. Our modeling considers the learning rate *α*_*t*_ and the received reward *r*_*t*_ to be in directly multiplicative relationship (6)—the two quantities are thus exchangeable. Inferring a (resp. dynamic) learning rate is thus akin to inferring a (resp. dynamic) reward *percept*, providing a relationship to latent value modeling of IRL [20]. Furthermore, the cost *and* the metric together define the geometry of gradient ascent learning flow. Our inferred positive, vector-valued learning rates can be interpreted as a personalized effective metric (per-parameter step size), placing our results within recent (natural) gradient ascent perspectives on neural learning dynamics [36, 37].

Looking ahead, individually inferred learning models can help in identifying neural correlates of learning, allowing principled comparisons across species, tasks, and labs. They can further be leveraged for personalized curriculum design that is also adaptive to internal states, enhancing animal training.

## Acknowledgments

The authors would like to thank Y. Helena Liu, Yoel Sanchez Araujo, Lenca Cuturela, and Nathaniel Daw for helpful discussions. VG was supported the Porter Ogden Jacobus Fellowship at Princeton University, and by doctoral scholarships from the Natural Sciences and Engineering Research Council of Canada (NSERC) and the Fonds de recherche du Québec – Nature et technologies (FRQNT). JWP was supported by grants from the Simons Collaboration on the Global Brain (SCGB AWD543027), the NIH BRAIN initiative (9R01DA056404-04), an NIH R01 (NIH 1R01EY033064), and a U19 NIH-NINDS BRAIN Initiative Award (U19NS104648).

## A Data & pre-processing

### Data source and links

The data is pulled from the International Brain Laboratory (IBL) via the ONE API on OpenAlyx: https://openalyx.internationalbrainlab.org (docs: https://int-brain-lab.github.io/ONE/). IBL project site: https://www.internationalbrainlab.org.

### Subject/session selection

- We identify mice that progressed to the `biasedChoiceWorld` training protocol (“trained” cohort) and collect their sessions for the requested `protocol` (e.g., training or `biasedChoiceWorld`). The no_curriculum variant filters sessions by the tag `2024_Q3_Pan_Vazquez_et_al`.
- We keep only sessions that contain all required trial fields: {contrastLeft, contrastRight, choice, probabilityLeft, feedbackType, rewardVolume}.

If any session for a subject fails to load or is malformed, that entire subject is excluded (logged as an “error animal”).

### Behavioral and trial recoding

We apply some transformations to the loaded data.

- Missing contrasts are set to 0 (NaNs only occur in contrast entries).
- We convert the raw wheel choice to *Y* ∈ {−1, 0, +1} (left, no-go, right) using the task’s sign convention; apply a sign flip so that rightward is +1.
- *No-decision handling:* trials with no decision (*Y* = 0) are recoded to *Y* = +1 (right).
- We map `feedbackType` from {−1, +1} to {0, 1}.
- Sanity check: on trials with nonzero contrast, assert that `feedbackType` = **1**[choice = correct side].

### Design matrix *X* (regressors)

- Supported regressors include: stimIntensity, contrastLeft, contrastRight, correctSide, previousChoice, previousRewarded.
- We use a **tanh transform**, following [16]. For a contrast *c*, define tanh 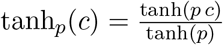 and we use *p* = 5. We apply it to `contrastLeft` and `contrastRight`.
- `correctSide: +1 if contrastRight > contrastLeft, else −1`.
- `previousChoice: previous` trial’s *Y* (first trial padded with 0).
- `previousRewarded: previous trial’s correctSide` (first trial padded with 0).

## B Modeling

### B.1 Probabilistic models definitions

We consider various models in this work:

1. **PsyTrack**: We consider the PsyTrack model from [16], which is obtained by setting *α* = 0 in eq. (2). For completeness, the generative model reads as

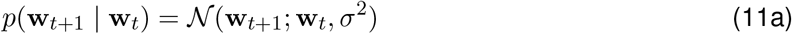

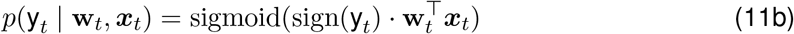

with *σ* replaced by *σ*_day_ on trials *t* representing a switching between training days. We refer the reader to [16] for a more thorough explanation of the model. The parameter set in this model is *θ* = {*σ, σ*_day_}, which we optimize by maximizing the marginal likelihood *p*(**y**_1:*T*_ | *θ*), marginalizing over weight **w**_1:*T*_ trajectories.
2. **TDRL**: This model refers to an agent within a POMDP learning by TDRL as per [28]. We outline it below for completeness but refer the reader to [28] for a more thorough explanation of the model. The policy can be described as a softmax policy with temperature *β >* 0

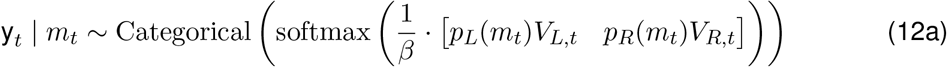

given the expected-reward “*Q*” values for each side formed from left and right values {*V*_*L,t*_, *V*_*R,t*_} and the posterior belief distribution

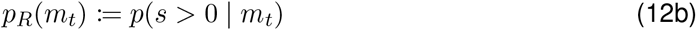

about the true stimulus intensity *s* given a noisy percept 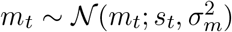. Once the decision is made and a reward is received, the values for the choice *c* = y_*t*_ ∈ {*L, R*} are updated using the reward prediction error as

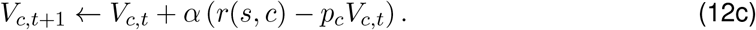

Finally, we relax these dynamics to be probabilistic, with additive Gaussian noise added to each values per trial. The parameter set in this model is *θ* = {*σ*_*m*_, *σ, σ*_day_, *β, α*}, which we optimize in the same fashion as all other models by maximizing the marginal likelihood *p*(**y**_1:*T*_ | *θ*), marginalizing over percept **m**_1:*T*_ and value {**V**_*L*,1:*T*_, **V**_*R*,1:*T*_ } trajectories.

### B.2 Relationship between TDRL action-value and GLM policy-gradient models

#### B.2.1 Equivalence in policy between action-value models and GLMs

Our aim is to establish a correspondence between the TDRL model in eq. (12) and our GLM setting, focusing on the resulting policies.

Consider the policy *p*(y_*t*_ = *R* | *s*_*t*_), the probability of decision y_*t*_ = *R* given the stimulus intensity *s*_*t*_ on trial *t*. As a Bernoulli draw, it is equal to 𝔼 [y |*s*] the expected value of the choice y ∈ {0, 1} in response to the stimulus intensity *s*. Let us omit the subscripts *t*. We have

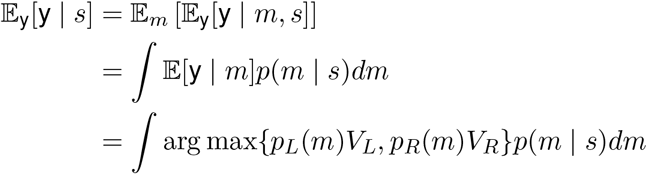

Where 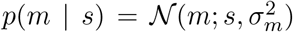. Note that we assume a uniform prior *p*(*s*) following [28], thus conversely 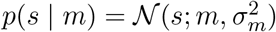.

Let us start with the simple case where *V*_*R*_ = *V*_*L*_ = 1 and *σ* = 1. We see that if 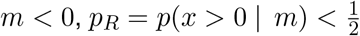, since the mean is negative, thus *p*_*L*_ *> p*_*R*_ and 𝔼 [y |*m*] = 0. Similarly for *m >* 0, 𝔼 [y |*m*] = 1. This helps us rewrite the above integral to a simpler one with limited support:

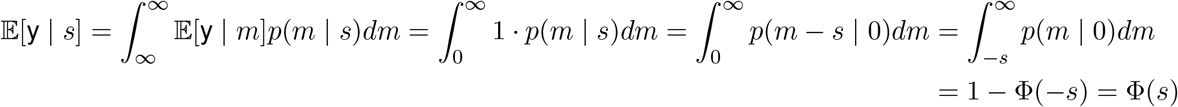

which shows that the expected choice follows the CDF Φ of the standard normal distribution. The following proposition details this idea for general {*V*_*R*_, *V*_*L*_}.

##### Proposition B.1

*For the TDRL model of eqs. (12) given stimulus intensity s* ∈ [−1, 1], *percept distribution* 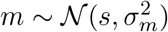 *and “argmax” decision rule with values* {*V*_*R*_, *V*_*L*_}, *the policy follows*

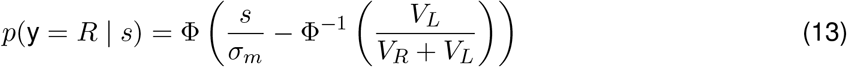

*for* Φ *the CDF of the standard Gaussian*.

*Proof*. For general *V*_*R*_, *V*_*L*_ and *σ >* 0, we can follow the same procedure of finding the transition point where *p*_*R*_ *> p*_*L*_, and rewriting the bounds of the integral. We have

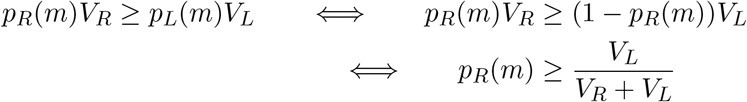

and furthermore

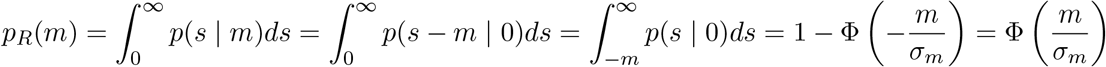

so the transition point follows

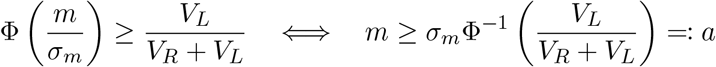

Plugging this back into our expected choice integral, we get

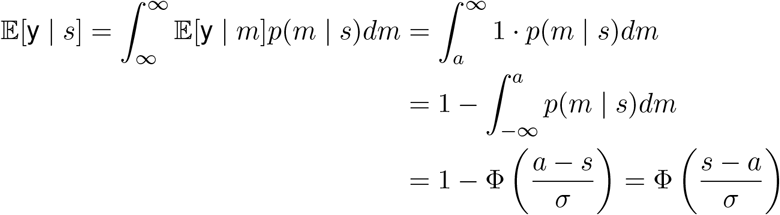

as desired. □

The function Φ is a “sigmoidal” function and closely approximates the sigmoid of the logit link function. One could have easily considered Φ for a *probit* link function in the GLM, which would have made the relationship exact, considering 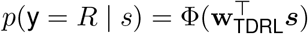 with the linear model

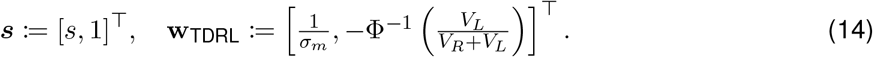

Staying with a logit link function, one can further use that 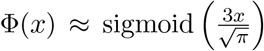 makes for a close approximation to the probit link.

We can interpret from (13) that the smaller the perceptual noise *σ*_*m*_ the stronger the gain of the sigmoid is, and the values {*V*_*L*_, *V*_*R*_} dictate the bias through their normalized difference. As a sanity check, we see that if *V*_*R*_ = *V*_*L*_, the transition point, i.e. the bias, satisfies *a* = 0.

#### B.2.2 Confidence-guided updating in policy-gradient learning

Prior work [28] emphasized the prevalence of confidence-guided updating during learning in perceptual decision-making experiments, manifesting as a bias in choice behavior following difficult trials. While these findings are introduced using the TDRL model described above, the primary contributing factor to this phenomenon is the incorporation of decision-confidence in the value updating itself. This is shared by the policy gradient learning methods considered in this work, as one can see from the multiplicative term dependent on *p*(y_*t*_ | **w**_*t*_, ***x***_*t*_) in eq. (6). We further show in Fig. 5 the scalar change in policy Δ*p*,

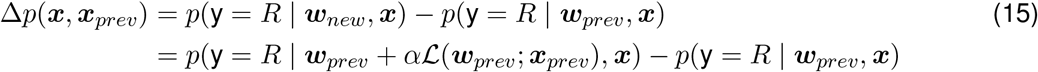

**Figure 5:**
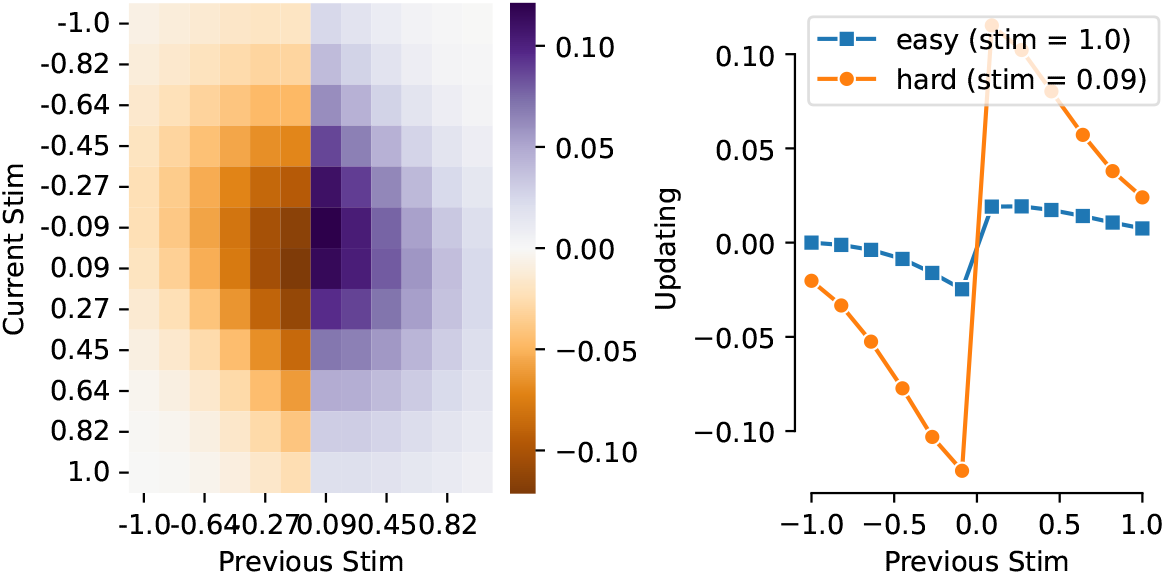
Confidence guided updating in policy-gradient learning. (**left**) heatmap of Δ*p*(***x, x***_*prev*_). (**right**) Δ*p* as a function of ***x***_*prev*_ for easy *s* = 1.0 and hard *s* = 0.09 current stimulus intensities.

**Figure 6:**
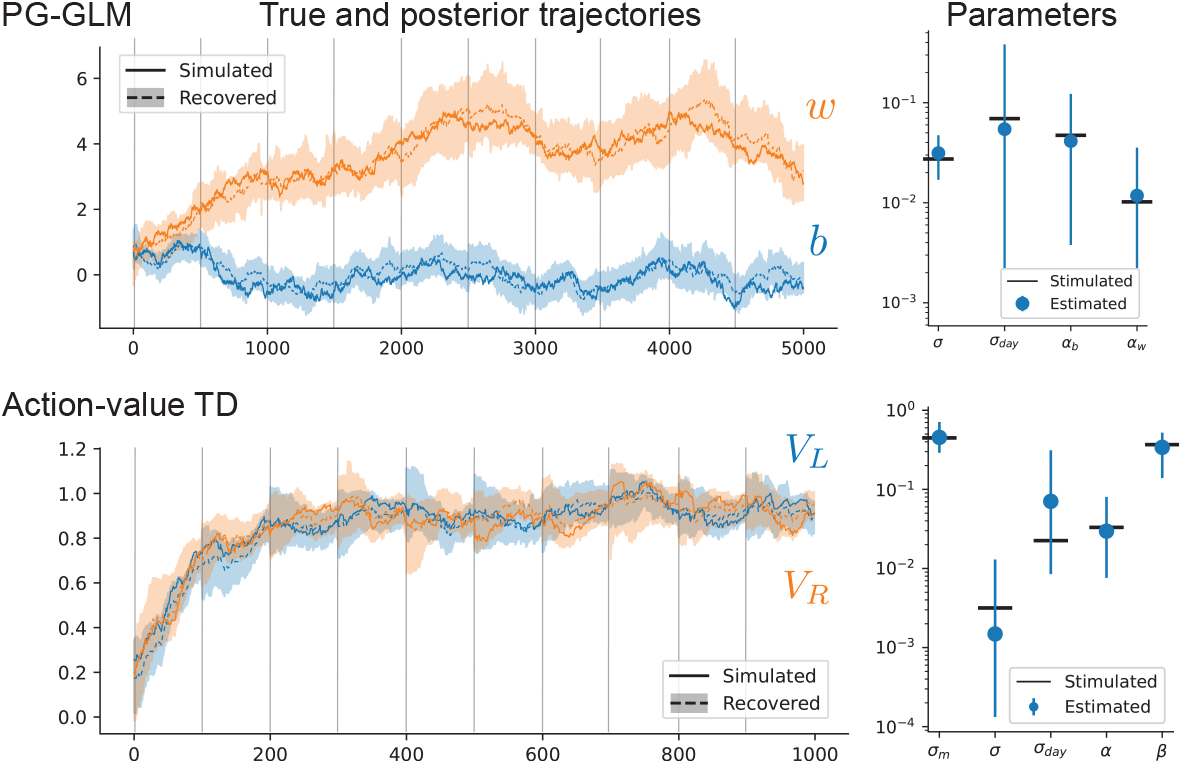
Recovery on simulated data. (**left**) Simulated trajectories (parameters on the right) of weight and biases updated under the vector PG model (top) and left-right values for the TDRL model (bottom) are faithfully recovered by the particle filtering algorithm. The error bars indicated one standard deviation, estimated from the particles. Faint lines correspond to simulated “sessions” days. (**right**) Parameter recovery from our MLL optimization procedure, with error bars indicating the 95% confidence interval from our posterior estimate with MCMC.

at a current stimulus ***x*** = [1, *s*]^⊤^ and previous stimulus ***x***_*prev*_, simulated for a fixed ***w***_*prev*_ = [0, 3]^⊤^ and *α* = 1 and with ℒthe full policy-gradient update. We see that we replicate the confidence-guided behavior [28].

#### B.2.3 Learning dynamics

The TDRL model takes the animal’s perspective when making decisions and learning. Both are taken given *m*, the agent’s representation of the true stimuli *s*. As experimenters, we have access to *s* only—the animal has access to *m* only. In our attempt to bridge the model with the GLM, we must consider what is the change in values given a stimulus *s*, averaging over internal percept *m*, characterized by the distribution

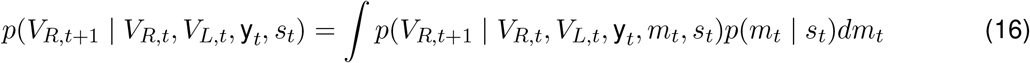

and similarly for *V*_*L*_.

Let us start by considering the mean of the distribution in (16), in *V*_*R*_, dropping subscripts *t* and denoting 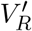 for *V*_*R,t*+1_. We have

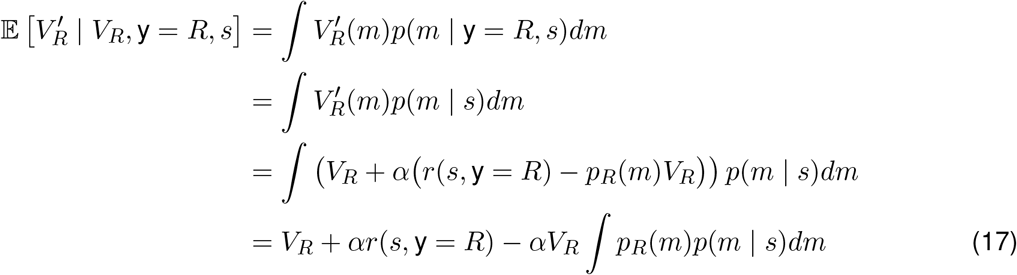

To solve this integral in (17), note that 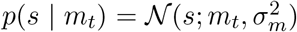 assuming uniform prior *p*(*s*), as done by [28]. As such,

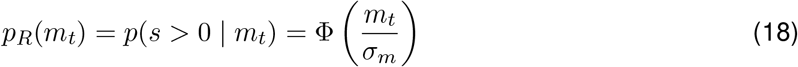

where Φ is the CDF of the standard Normal distribution. The integral of interested in (17) is thus the expected value of a Gaussian CDF under a Gaussian distribution, which is solvable. Consider 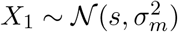 and 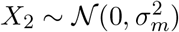. Then

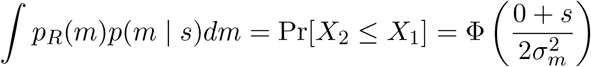

where we’ve used that *X*_2_ − *X*_1_ ∼ 𝒩 (−*s*, 2*σ*^2^). We’ve thus shown that

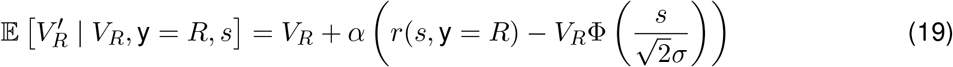

and similarly,

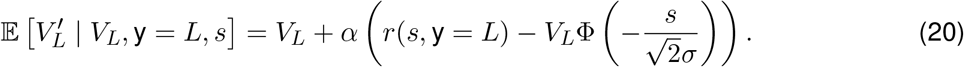

Next, we consider the second moment 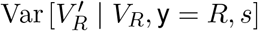 of this distribution. We can approximate it using the Taylor expansion

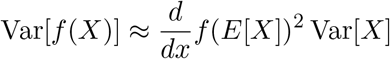

for *X* a random variable and *f* a scalar-valued function. With *f* (*m*) = *V*_*R*_ + *α* (*r*_*t*_ − *p*_*R*_(*m*)*V*_*R*_), we have

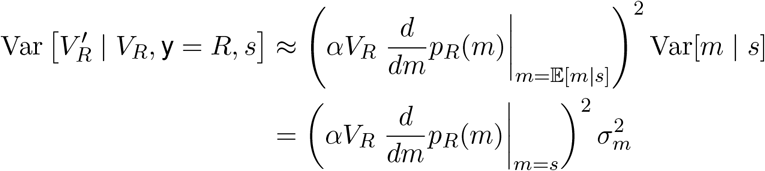

and

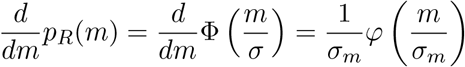

where *φ* is the PDF of the standard Normal distribution. Taken together, we have

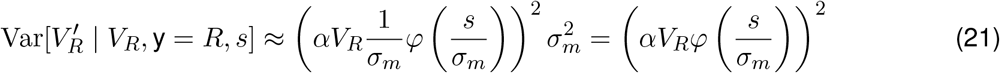

and similarly

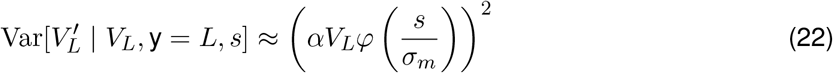

using the symmetry of *φ* about *s* = 0.

These two moment calculations provide the Gaussian approximation to the distribution

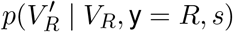

such that

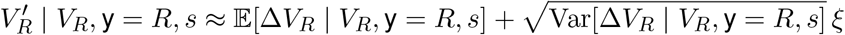

where *ξ* ∼ 𝒩 (0, 1). Similar derivations throughout hold for *V*_*L*_.

Taking it all together, given a choice *c* = y_*t*_, the change in values conditioned on the stimulus percept *m*_*t*_ is given by our model as

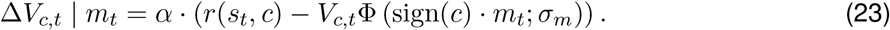

We’ve shown that the change in values conditioned only on the stimulus intensity *s*_*t*_, averaging over percepts *m*_*t*_, is approximated up to its first two moments as

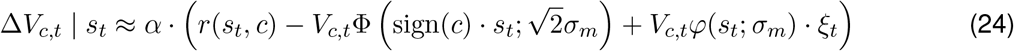

for *ξ*_*t*_ ∼ 𝒩 (0, 1), where Φ (*x*; *σ*) and *φ* (*x*; *σ*) are respectively the CDF and PDF of 𝒩 (*x*; 0, *σ*^2^).

We’ve thus shown that while the TDRL and PG-GLM are equivalent in policy, they differ in how their respective policy weights evolve. Namely, if we approximate the TDRL as a latent update over only the decision-making weights, i.e. the values, then we observe that the perceptual noise manifests also in the noise scaling of the weights update. This update noise is accentuated for higher values of *V*_*c,t*_ and low stimulus intensity *s*_*t*_, and decays as *σ*_*m*_ → 0.

### C Implementation and numerical considerations

### C.1 Inference procedure

#### C.1.1 Parameter inference

Our parameter inference, i.e. fitting, procedure decomposes into the following steps: to infer a parameter set *θ* of a learning model *p*_*θ*_(y_1:*T*_ | ***x***_1:*T*_) for a specific single-animal trajectory y_1:*T*_ :

1. *Train-test decomposition*: Sample 10% of indices from {1, …, *T* } to be held-out, and mask (i.e. NaN) the decisions y_*t*_ at these time-steps. The result is a (corrupted) train trajectory 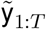 and the original trajectory y_1:*T*_ .
2. *Initialization*: Perform a grid search over a user-defined grid of parameters to obtain *θ*_init_. This step can be performed in parallel over parameter combinations. For nested models we initialize at parameters of sub-model and grid-search over remaining parameters.
3. Perform *maximum likelihood* inference

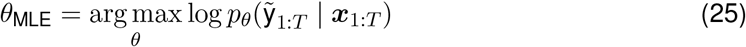

through gradient ascent initialized at *θ*_init_ until convergence. We detail how this objective function is estimated in the next subsection. We use the AMSGrad optimizer [38] with exponential decay (rate = 0.1, two cycles of decay) learning rate schedule initialized at *η* = 0.01.

#### C.1.2 Sampling-based estimation methods

##### Marginal log-likelihood with particle filtering

We report the full trajectory marginal log-likelihood as our primary metric

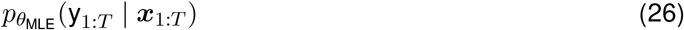

which re-introduces the held-out trials. This integral is high-dimensional and intractable, and we resort to sequential Monte Carlo (SMC) (also known as particle filtering) to compute it [24, 25]. We use *N* = 5000 particles for all computations reported.

##### Parameter posterior with MCMC

We use Metropolis Hastings to obtain samples from the posterior. As we posit a uniform prior over the parameters *θ*, we use the marginal likelihood *p*_*θ*_(y_1:*T*_ | ***x***_1:*T*_) as the target distribution proportional to the density of interest, the posterior *p*(*θ* | y_1:*T*_, ***x***_1:*T*_). We use a proposal distribution a Gaussian distribution over each parameter *θ*_*i*_ with uniform scale *σ*_proposal_ = 0.5. We run the algorithm for 100 iterations, with a 50 iteration burn-in, and 500 samples.

### C.2 Learning and noise decompositions

For a given animal *i*, consider

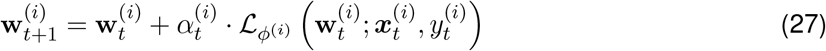

the noiseless learning dynamics, at time step *t* ∈ {1, …, *T* ^(*i*)^} from previous weights 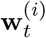, and animal-specific regressors 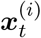 and decision 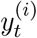 from the data, which define the *learning trajectory*. The learning rate 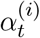 is fixed to the inferred parameter ***α***^(*i*)^ for the vector PG model, and set to the mean 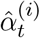 of the posterior 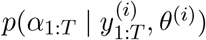 for the dynamic PG model.

Dropping the animal *i* superscripts henceforth, let us now detail the learning metrics we consider. Take ŵ_1:*T*_ the posterior mean, and let 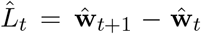 be the corresponding update at trial *t*. Let *L*_*t*_ = *α* · ℒ (**w**_*t*_; ***x***_*t*_, y_*t*_) be the learning update from previous weights **w**_*t*_, regressors ***x***_*t*_ and sampled decision y_*t*_. The main metric L we consider is the *learning fraction*

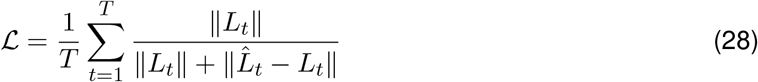

which is the average, over trials, of the fraction between ∥*L*_*t*_∥ and the norm of the residual 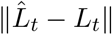 (Fig. 7**d**).

We considered also the following alternate metrics, for which we plot in Fig. 7**d**-**f** their values over animals.

1. The *projection fraction* (Fig. 7**e**) is the average of the absolute projection ratio between 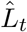 and *L*_*t*_.

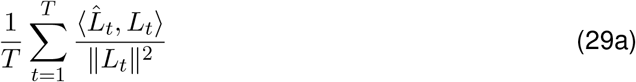
2. The *cosine similarity* (Fig. 7**f**) metric is the average of the cosine similarity between 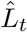 and *L*_*t*_

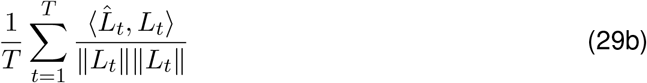
3. The *learning fraction #2* (Fig. 7**g**) is the metric considered in Ref. [22], the average of the squared cosine similarities

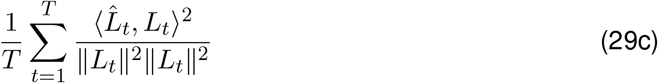

### C.3 Weight non-identifiability in GLM likelihood

Here we discuss weight **w** identifiability considerations for the Bernoulli GLM policy model considered so far. We define weights **w**_1_ and **w**_2_ to be *non-identifiable* on the basis of a function *f*, like the likelihood, if *f* (**w**_1_) = *f* (**w**_2_).

Consider the Bernoulli GLM policy studied so far, with left and right stimulus intensities *x*_*L*_ and *x*_*R*_ vectorized into ***x*** = [*x*_*L*_, *x*_*R*_, 1] and weights **w** = [*w*_*L*_, *w*_*R*_, *b*],

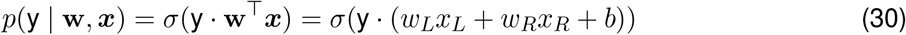

for y ∈ {−1, 1}. Now, consider the following perturbation

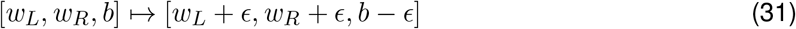

for *ϵ >* 0. We will show that this kind of perturbation has minimal to no effect on the likelihood, but can have a substantial effect on the evolution of the weights. Indeed, let *f* (*ϵ*; **w, *x***) be the likelihood of y = *R* with this perturbation

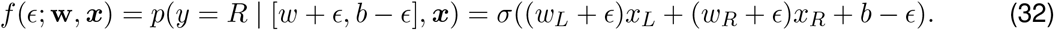

We can examine the influence of *ϵ* on *f* through it’s impact on the derivative. It is easy to see that

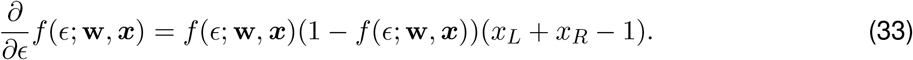

from which we can conclude a few things

- If either 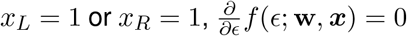 meaning that this kind of perturbation would have no impact on the log-lik;
- As *ϵ* increases, 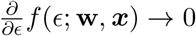 meaning it has less and less of an impact on the log-lik;

see Fig. 8**a** for a heatmap of 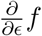 over {*s, ϵ*} for *s* = *x*_*R*_ − *x*_*L*_ the stimulus intensity.

This creates a “weak” form of non-identifiability where substantial changes in the weights, or parameters, will have small (non-zero) changes in the log-likelihood, becoming strict at the boundary |*s*| = 1 We noticed this behavior in Psytrack – see an example below in Fig. 8**c**. We elect thus to fix the bias term to 0 until contrast of sufficiently low-levels have been observed in the data.

## D Supplementary results

**Figure 7:**
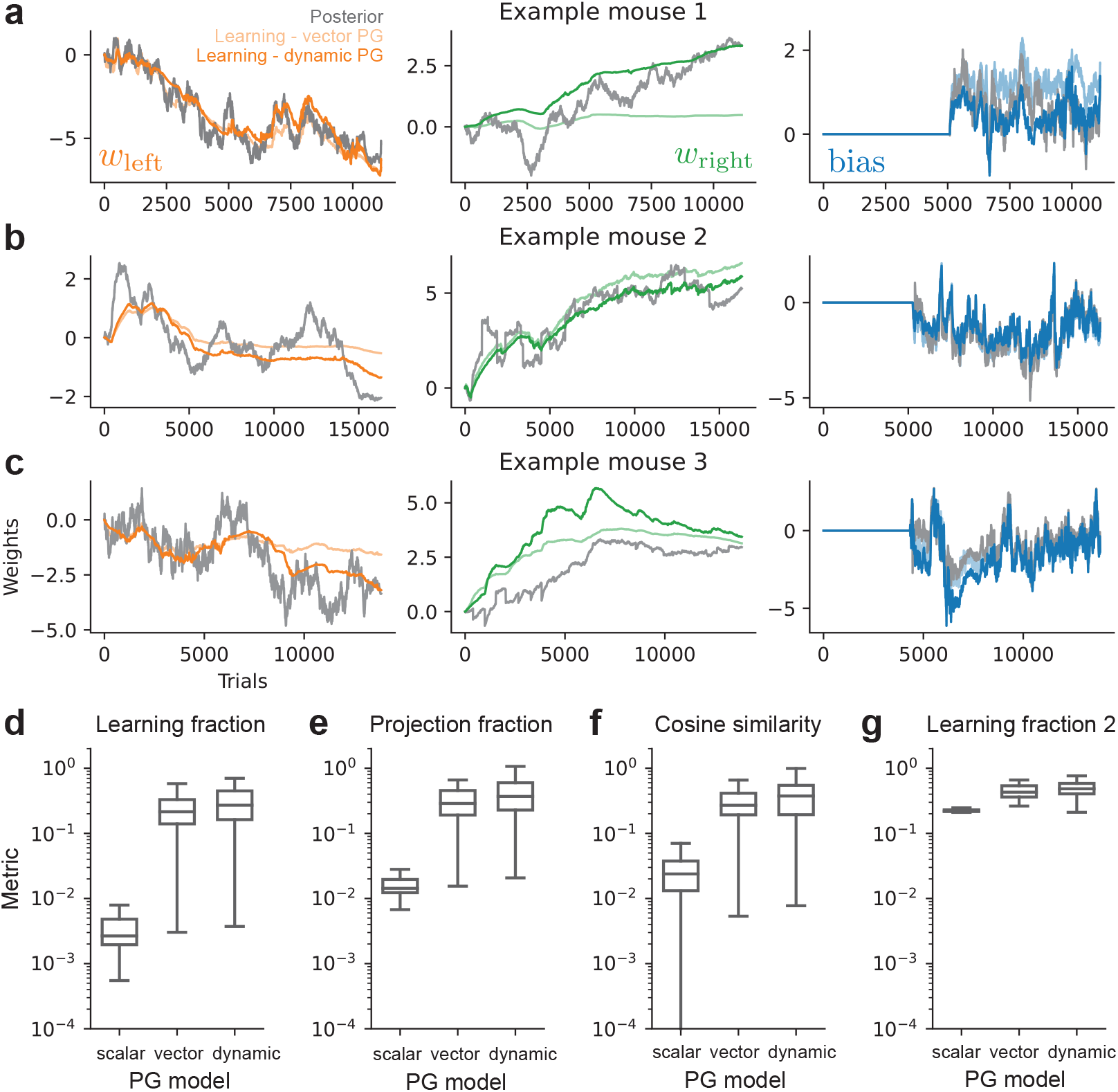
Learning and noise decomposition supplemental. (**a**-**c**) Learning component trajectories for both vector and dynamic PG models for all three mice considered, per regressor component, overlaid over the posterior mean ŵ_1:*T*_ trajectory under the vector PG model (similar to dynamic PG). The learning component of the dynamic PG is broadly more aligned with the inferred posterior. (**d**-**g**) Metric of learning and noise decomposition, per PG model, with boxplots indicating distributions of individual metric scores over animals. Higher is better. See §C.2 for more details—we used the learning fraction in (**d**) in the main text.

**Figure 8:**
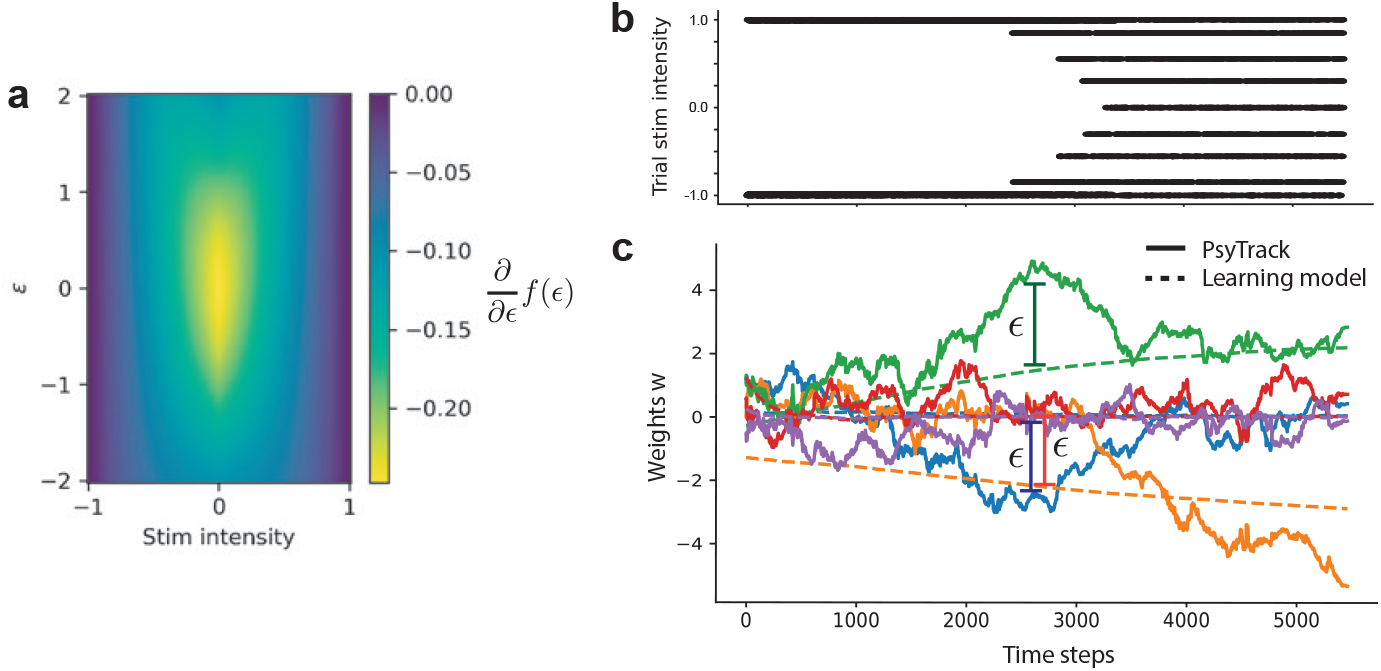
(**a**) Heatmap of 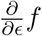 over {*s, ϵ*}. The gradient vanishes (i.e., non-identifiability) near |*s*| = 1. (**b**) Stimuli sequence, over trials, presented to an example animal. (**c**) Inferred mean posterior weight trajectories under PsyTrack and REINFORCE, without bias correction. The *ϵ* deviations are visible.

**Figure 9:**
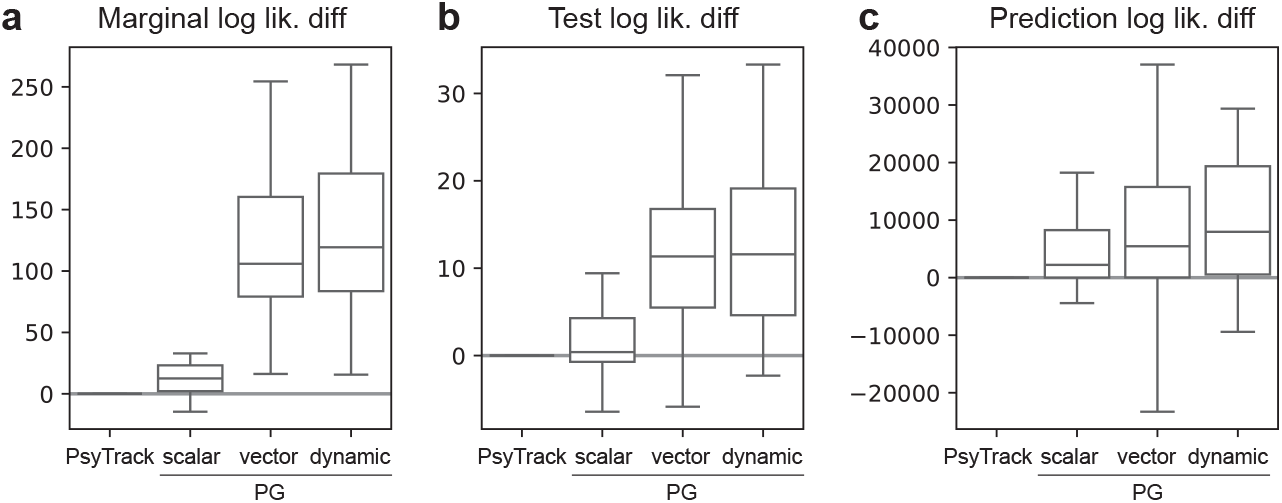
Performance metrics considered, across PG models, with boxplots indicating distributions over animals, shown as difference w.r.t. to PsyTrack [16] (higher is better). The marginal log-lik. is defined on the entire trajectory, whereas the test log-lik is the sum of the filtering log-liks. on over the 10% held-out trials. The prediction log-lik. is an alternate metric obtained by simulating trajectories under the inferred generative model, that is *sampling* decisions y_*t*_ at each time step, and using the resulting emission likelihoods evaluated at the *true* (data) decisions.

**Figure 10:**
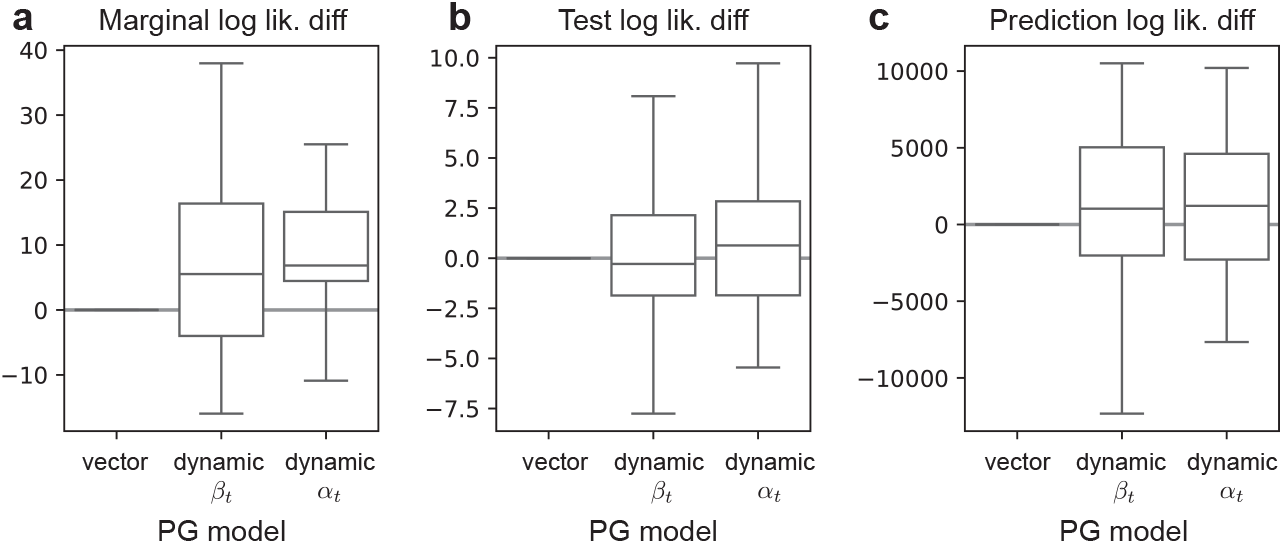
Performance metrics across PG models. We compare the vector PG model to the dynamic learning rate variant considered in the main text (§2.4), as well as a dynamic PG model with dynamic vector baseline ***β***_*t*_, defined similarly to the dynamic learning rate model as to strictly generalize the vector PG model. We see that the dynamic *α*_*t*_ model outperformed the two others.

